# Characterizing the Behavioral and Neuroendocrine Features of Susceptibility and Resilience to Social Stress

**DOI:** 10.1101/2021.12.13.472392

**Authors:** Dalia Murra, Kathryn L. Hilde, Anne Fitzpatrick, Pamela M. Maras, Stanley J. Watson, Huda Akil

## Abstract

Evaluating and coping with stressful social events as they unfold is a critical strategy in overcoming them without long-lasting detrimental effects. Individuals display a wide range of responses to stress, which can manifest in a variety of outcomes for the brain as well as subsequent behavior. Chronic Social Defeat Stress (CSDS) in mice has been widely used to model individual variation following a social stressor. Following a course of repeated intermittent psychological and physical stress, mice diverge into separate populations of social reactivity: resilient (socially interactive) and susceptible (socially avoidant) animals. A rich body of work reveals distinct neurobiological and behavioral consequences of this experience that map onto the resilient and susceptible groups. However, the range of factors that emerge over the course of defeat have not been fully described. Therefore, in the current study, we focused on characterizing behavioral, physiological, and neuroendocrine profiles of mice in three separate phases: before, during, and following CSDS. We found that following CSDS, traditional read-outs of anxiety-like and depression-like behaviors do not map on to the resilient and susceptible groups. By contrast, behavioral coping strategies used during the initial social stress encounter better predict which mice will eventually become resilient or susceptible. In particular, mice that will emerge as susceptible display greater escape behavior on Day 1 of social defeat than those that will emerge as resilient, indicating early differences in coping mechanisms used between the two groups. We further show that the social avoidance phenotype in susceptible mice is specific to the aggressor strain and does not generalize to conspecifics or other strains, indicating that there may be features of threat discrimination that are specific to the susceptible mice. Our findings suggest that there are costs and benefits to both the resilient and susceptible outcomes, reflected in their ability to cope and adapt to the social stressor.

## INTRODUCTION

For humans, social interactions can be a source of support or stress. The nature of interpersonal relationships and interactions plays a crucial role in the development of an affective state. Whereas positive social relationships can strengthen mental and physical well-being, negative social relationships have the propensity to induce highly stressful and harmful environments. During social stress, different individuals rely on various coping styles or strategies that may prove either adaptive or maladaptive in the long term (Billings and Moos, 1984; Connor-Smith and Compas, 2002; Wood and Bhatnagar, 2015).

In rodents, social defeat studies provide a naturalistic dominance model that uses agonistic encounters as a trigger, and social coping as a dependent measure (Hollis and Kabbaj, 2014). An agonistic encounter is often characterized by intense, aggressive interactions among animals of the same species that ultimately leads to individuals emerging as either ‘winners’ or ‘losers’ following the stress (Kollack-Walker et al., 1999; Kabbaj et al., 2001; Hammels et al., 2015). In mice, the Chronic Social Defeat Stress (CSDS) model relies on a paradigm where an experimental mouse is exposed physically and psychosocially to a more aggressive mouse from a different strain. Following the end of the chronic stress, the defeated animals diverge into two separate populations of social reactivity: a “resilient” group that is socially interactive and a “susceptible” group that is socially avoidant (Golden et al., 2011).

Defeated mice are termed ‘resilient’ and ‘susceptible’ based on their social behavior after stress. Understanding when and how these reactivities arise is critical in identifying predictive factors that set the course for vulnerability. For example, baseline differences in anxiety-like behaviors (Milic et al., 2021), as well as neuroanatomy and physiology (Nasca et al., 2019), have been shown to predict the social avoidance outcome. Additionally, social hierarchies established in the homecage, well before the stress, predict the resilient and susceptible outcome. This is particularly interesting, as it suggests that there may be features of the defeat (e.g., dominance) that reflect intrinsic sociability. During defeat encounters, mice may engage in a variety of active and passive coping behaviors, such as fight, escape, and freezing, as they react to the social stressor (McLaughlin et al., 2006). Also involved in the interaction is the ability to distinguish between threatening and non-threatening social cues, such as the environmental context and aggressor (Ayash et al., 2020). Observing the features of these adaptive and reactive strategies that resilient and susceptible mice use *during* stress may allow us to identify additional predictors of social outcome and paint a more cohesive picture of what constitutes vulnerability.

The behavioral and physiological components of the stress response are deeply intertwined; therefore, it is important to understand how interactions between these traits map onto different social outcomes following social defeat. The primary driver of the stress response is the hypothalamic-pituitary-adrenal (HPA) axis. When activated, a cascade of events leads to the adrenal synthesis and release of corticosteroid hormones— specifically, corticosterone (CORT) in rodents and cortisol in humans—targeting numerous organs, including the brain (Akil, 2005). Through this neuroendocrine axis, stressful experiences can trigger a wave of physiological consequences, including alterations in the HPA axis itself, as well as changes in body weight regulation and pain sensitivity (Abdallah and Geha, 2017; Butler and Finn, 2009; Iio et al., 2014; Jeong et al., 2013). In humans, affective disorders, including major depressive disorder, have been associated with shifts in circadian cortisol rhythms (Adam et al., 2017) and an initial increase in cortisol in response to a social stress (Adam et al., 2017; Young et al., 2000).

To understand the specific factors that may lead an animal to display features of resilience or susceptibility to social stress, we conducted a series of careful studies examining behavioral and endocrine variables across the CSDS paradigm. We examined mice before (pre-), during, and after (post-) CSDS (**Fig. 1A**), generating a detailed temporal characterization of resilient and susceptible mice and allowing identification of key variables that predict phenotype divergence. Importantly, our results suggest that social reactivity to CSDS can be visualized during the defeat itself via inherent differences in coping mechanisms and that social reactivity is specific to the social context from a behavioral, neuroendocrine, and physiological perspective.

**Figure 1:**
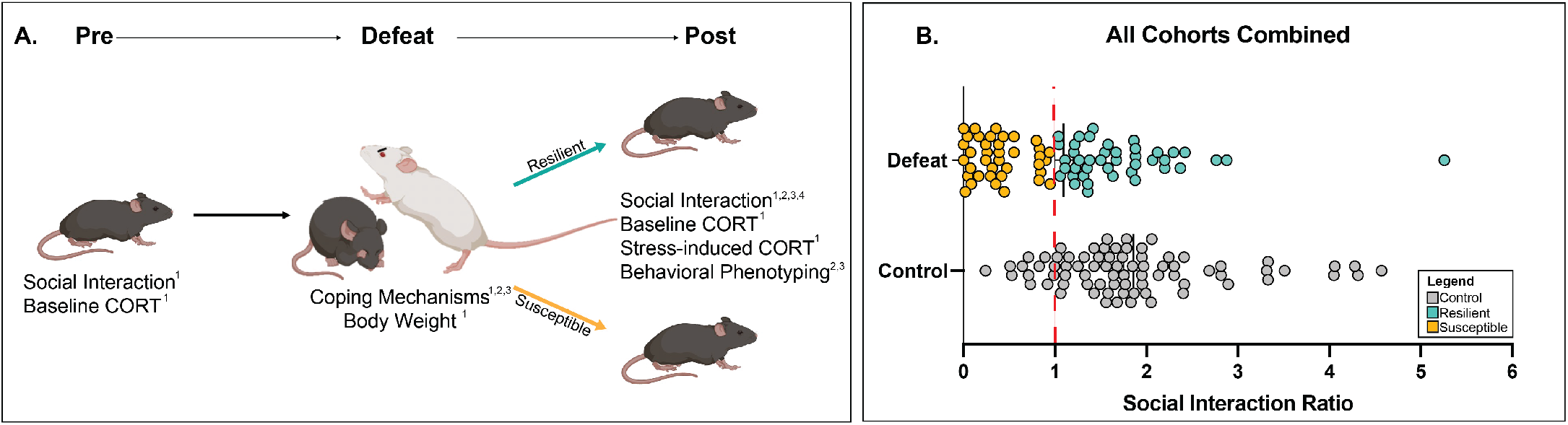
Experimental Schematic: Behavioral and endocrine profiling following chronic social defeat stress (CSDS) **1A.** Schematic timeline of the 3 timepoints (pre-, during, and post-defeat) used for behavioral phenotyping and endocrine profiling. Each timepoint includes the overall category of experiments conducted. Superscripts indicate the cohorts of mice used for each assay. **1B**. Social Interaction (SI) ratios of all 4 cohorts of mice combined. Defeat (*n*=80; *mean*=1.096, *sd*=+/− 0.877) and Control (*n*= 77; *mean*= 1.856, *sd*=+/−0.934). Fig. 1A created with BioRender. Colors: red dash= SI ratio of 1, yellow = susceptible, blue = resilient, and grey = control.

## METHODS

### Animals

All mice were purchased from Charles River Laboratories (Wilmington, MA) and allowed at least one week for habituation to the vivarium prior to use in experiments. Subjects were C57BL/6J male mice aged 8-11 weeks, housed 2-4 per cage upon arrival and up until the experiment start. Male CD1 retired breeders (3-6 months) were used as aggressors and social interaction stimulus mice. CD1 mice were singly housed one week prior to aggressor screening and the start of the experiment. Black Swiss mice (10-14 weeks) were used as stimulus animals in a subset of social interaction tests. Enrichment in the form of enviro-paks was provided throughout the study. The rooms were kept under constant temperature (25 ± 2°C) and with a 12 h light:12 h dark cycle starting at 7 AM. All behavioral tests were conducted under standard overhead lighting conditions unless otherwise noted. Mice were provided with mouse chow and tap water *ab libitum* and maintained in accordance with the *NIH Guidelines for the Care and Use of Laboratory Animals.* The University of Michigan Institute of Animal Use Care and Use Committee (IACUC) approved all animal protocols utilized.

### Chronic Social Defeat Stress (CSDS) and Control Rotation (CR)

CSDS was performed according to the previously described protocol (Golden et al., 2011). Briefly, this paradigm consisted of a C57BL/6J intruder male mouse repeatedly subjected to 5-minute bouts of physical defeat from a resident CD1 mouse. All CD1 mice were screened for aggression (defined as < 1-minute interval to aggressive behavior) prior to use in CSDS experiments. Following the defeat, the intruder mouse was placed across the resident CD1 in a (26.7×48.3×15.2 cm) cage with a perforated plexiglass divider between them for 24 hours to elicit the psychosocial aspect of the stress. Each day for ten days, the intruder mouse was rotated to a new CD1 resident to prevent habituation. Control mice were housed 2/cage across a plexiglass divider and rotated to a new conspecific each day for ten days. All CSDS and CR occurred between 9 AM-11 AM each day. Following CSDS or CR, experimental mice were singly housed in the vivarium with enviro-pak enrichment.

Defeat sessions were recorded with a video camera and manually scored by a blinded observer. The behaviors measured were total duration of engagement in the fight and total number of fights, escapes, and upright, forward, and crouch-back freezing during physical defeat bouts (Table 1). Duration measurements are reported in seconds, and all other behaviors are reported as a ratio of behavior measured/total behaviors. Three cohorts were recorded for this analysis, resulting in a total of *n*=22 resilient mice and *n*=26 susceptible mice.

**Table 1.** Summary table of behaviors scored during physical defeat bouts

| Behavior Scored | Description |
| --- | --- |
| Fight | Intruder mouse actively engages in fight |
| Escape | When approached, the intruder mouse jumps over the resident mouse |
| Upright Freeze | Intruder mouse freezes while on hind legs |
| Forward Freeze | Intruder mouse freezes while on all four legs, facing towards the resident |
| Crouch Back Freeze | Intruder mouse freezes with all four legs pulled in, facing away from resident |

**Table 2.** Testing conditions per experimental cohort

| Cohort | Group Size | Test Conditions | Testing Timelines |
| --- | --- | --- | --- |
| 1 | $n=20$ control, $n=16$ defeat | SI, Baseline CORT, FIT CORT, Body Weight | Figs. 2A & 5A |
| 2 | $n=20$ control, $n=15$ defeat | SI, OF, FST | Fig. 4A |
| 3 | $n=19$ control, $n=17$ defeat | SI, VF | Fig. 4A |
| 4a | $n=10$ control, $n=17$ defeat | SI w/ CD1, SI w/ Novel C57BL/6J | Fig. 6A |
| 4b | $n=8$ control, $n=15$ defeat | SI w/ CD1, SI w/ Black Swiss | Fig. 6A |

### Social Interaction Test (SI)

SI was performed according to a previously described protocol (Golden et al., 2011). SI tests were conducted under red light in a testing arena (31×31×30 cm), with a removable stimulus cage (10×10×10 cm) used to hold the social stimulus animal. There were two, 2.5-minute trials. During the first trial, the experimental mouse was allowed to explore the arena with an empty stimulus cage. The mouse was then removed from the arena, and a social partner (CD1, unless otherwise noted) was placed inside the stimulus cage. The experimental mouse was returned to the arena for another 2.5 minutes. Each session was recorded and exploration of the whole arena, as well as the exploration of the cage/interaction zone was scored using automatic tracking software Noldus Ethovision XT software (Noldus Information Technology, Leesburg, VA). The SI Ratio was calculated as time spent in the interaction zone with CD1 present/not present. A social interaction score of ≤ 1 indicates susceptible, while > 1 indicates resilient (Assessed in Cohorts 1-4).

### Body Weight (BW)

Mice were weighed on Day 1 and again on Day 10 of the CR or CSDS. Change in body weight was measured as Day 10 (g) – Day 1 (g). (Assessed in Cohort 1).

### Corticosterone (CORT) Measurements

#### AM/PM Measurements

To assess the circadian rhythm of circulating CORT, AM and PM CORT were assayed three days before and following CSDS or CR. Blood was drawn two hours before lights on and one hour following lights off in the mouse vivarium via tail vein under red light using Sarstedt Inc. MICROVETTE CB300 EDTA tubes (Ref#16.444.100). Blood was centrifuged, and plasma was stored at −80°C until used for CORT analysis with the Arbor Assays CORT ELISA kit (Cat#K014-H1). (Assessed in Cohort 1).

#### Forced Interaction Test (FIT)

After CSDS or CR and one day following AM/PM blood draws, mice were placed across from a novel mouse in a clean CSDS cage, separated by a perforated plexiglass wall for 40 minutes to simulate both SI test and CR/CSDS conditions. No physical interaction was allowed, but sensory interaction was allowed through the perforated divider. Immediately upon the conclusion of this 40-minute test, mice were sacrificed, trunk blood was collected in 1mL tubes with 25 μL of 0.05M EDTA and centrifuged. Plasma was then stored at −80°C until used for CORT analysis with the Arbor Assays CORT ELISA kit (Cat#K014-H1). (Assessed in Cohort 1)

### Open Field (OF)

After CSDS or CR and one day following SI testing, mice were placed in an arena of 31×31×30 cm in size made of white plexiglass. Mice were allowed to explore the open field for 10 minutes. Time spent in the center was scored by Noldus Ethovision XT software. (Assessed in Cohort 2).

### Forced Swim Test (FST)

After CSDS or CR and two days following SI testing, mice were recorded from above and placed in a tall container (30×10 cm) filled within 2 inches from the top with 25 +/− 2°C water for 6 minutes. Total time spent immobile was hand-scored using a timer by an observer blind to the groups. Time spent immobile was defined as the least amount of movement to stay afloat (one hind-paw peddling or total immobility). (Assessed in Cohort 2).

### Von Frey (VF)

After CSDS or CR and one day after SI testing, each mouse was placed in a Plexiglas box atop a mesh platform and allowed to habituate for 30 minutes each day for three days. On the third day, mice were tested. Von Frey filaments of various forces starting from lowest to highest (0.008(g) - 2.0(g) Aesthesio Precise Tactile Sensory Evaluator Item#415000-20C)) were placed on the plantar surface of the hind paw (of each foot) until the fiber bends or the mouse withdrew its paw from the stimulus. Each foot was tested twice for force necessary to cause a withdrawal response and then averaged for the final score. (Assessed in Cohort 3).

### Cohort Breakdown

Four cohorts of mice were used to explore the above behaviors and physiological differences. Breakdown of group sizes and testing conditions are as follows:

#### During Stress

Cohorts 1, 3, and 4a were videotaped and scored for ‘During Stress’ Scoring (**Fig. 3A**).

### Statistics

Graphpad Prism and R (version 4.0.2) were used for figure design and statistical analyses. In R, *pastecs, car, afex, nlme, and emmeans* packages were used for statistical analyses (Fox and Weisberg, 2019; Grosjean and Ibanez, 2018; Lenth, 2020; Pinhero et al., 2020; Singmann et al., 2016). To determine main effects, one-way ANOVAs were evaluated with the *car* package, (aov function). To determine main and interaction effects with multiple time points, 2×2 ANOVAs were evaluated using the *afex* package (aov_ez function), with between factor = Group (control, resilient, susceptible) and within factor= time or tester strain. To control for cohort effects for the during stress behaviors, we added ‘cohort’ as a covariate to the 2×2 ANOVA. Post-hoc tests were conducted with the *emmeans* package (emmeans and pairs function), using a Tukey-HSD test. Mann-Whitney T-test was used for single comparisons among control and defeat groups. Pearson correlations were conducted via the cor.test function to assess linear relationships between outcome variables. Multi-level model (MLM) for baseline CORT data was evaluated using the *nlme* package (lme function) with estimates optimized using Restricted Maximum Likelihood (REML) and summarized to produce individual coefficients, approximate standard errors, and respective *p*-values for each level of the fixed effects variables. MLM was used to examine datasets that contain both repeated measures and missing data. There were missing data points for baseline AM/PM CORT samples due to not collecting enough blood from the tail vein (data point= blood collection time point--9 data points missing in controls, 7 data points missing in defeated mice). All significance thresholds were set at *p*≤0.05. Detailed statistics are included in the Supplemental Tables.

## RESULTS

### Characterizing the Resilient-Susceptible Outcome

We first examined social interaction behavior following ten days of chronic social defeat stress (CSDS) or control rotation (CR). Across four cohorts, control mice demonstrated a preference for social interaction, with a mean SI Ratio of 1.856 (*sd*=+/−0.934). In contrast, animals that experienced social defeat displayed a significant reduction in SI Ratios, with a mean of 1.096 (*sd*=+/−0.877) (**Fig. 1B**; *p*<0.001, *Mann-Whitney U*=1600). Defeat increased the proportion of socially avoidant mice (SI Ratio ≤1) from 16% (controls) to 48% (defeat). Notably, defeated mice could now be grouped according to two distinct response patterns: socially avoidant (susceptible) and socially interactive (resilient).

To understand this shift in population distribution due to defeat, we asked how the final susceptible and resilient features evolve longitudinally. We characterized defeated mice by observing features of behavioral, physiological, and neuroendocrine read-outs before (pre-), during, and after (post-) CSDS (**Fig. 1A**). The information gathered was used to identify which time(s) and which behavior(s) can predict the divergence of resilient and susceptible mice. This allowed us to ask whether the social avoidance phenotype is apparent early on, or whether it emerges due to the full extent of the repeated stress.

### Pre-Defeat: SI Ratios and CORT Levels Were Not Predictive of Future Reactivity to CSDS

#### Pre-CSDS SI ratios did not correlate with post-CSDS SI ratios

Given that the most robust finding following CSDS is the split of resilient and susceptible behaviors in the SI test, we asked whether this social behavior is a stable trait that 1) precedes stress and 2) could be used to predict resilient and susceptible groups. To that end, we conducted SI testing one day before and one day after CSDS and CR in the same animals (**Fig. 2A)**. There was no correlation between SI ratios pre-or post-CR in control mice, nor pre- or post-CSDS in defeated mice (Control: **Fig. 2B, Supp. Table 2;** *p*>0.05, Defeat: **Fig 2C, Supp. Table 2;** *p*>0.05). These data demonstrate that the development of vulnerability to CSDS cannot be predicted by social behavior before experiencing defeat.

**Figure 2:**
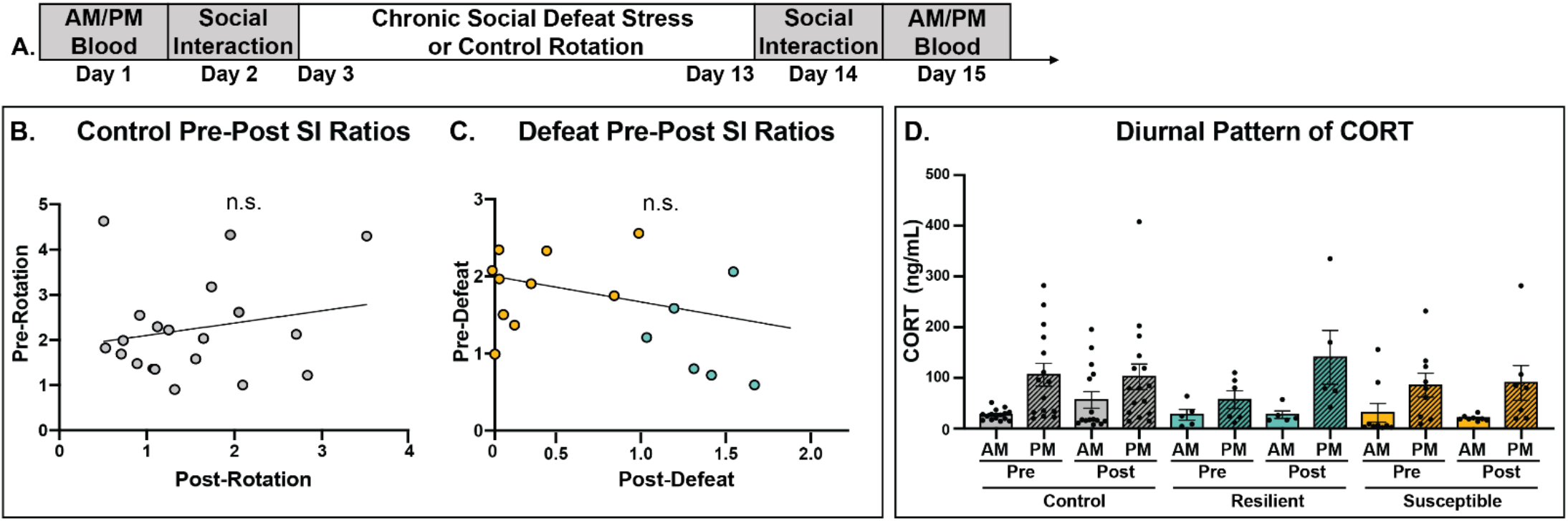
Pre-stress: Basal social interaction and corticosterone levels did not predict resilience or susceptibility post social defeat. **2A.** Schematic timeline of pre-post SI test and AM/PM corticosterone (CORT) sampling. **2B.** In control mice, SI Ratios displayed before the rotation did not correlate with SI ratios measured after the rotation. **2C.** In defeated mice, SI ratios measured before defeat did not correlate with SI ratios displayed after defeat. **2D.** CORT ELISA showing diurnal patterns of CORT before and after CSDS/CR. For all groups, the diurnal CORT patterns remained stable throughout the experiment, with higher PM CORT. For either pre- or post-CSDS/CR timepoints, there were no differences in CORT levels between groups. Significance Codes: n.s. = not significant. Colors: grey = control, blue = resilient, yellow = susceptible.

#### Social reactivity outcome was not predicted by basal diurnal CORT rhythms, either before or after CSDS

We then wanted to assess if baseline neuroendocrine differences predicted resiliency or susceptibility. There is a distinct diurnal pattern of CORT in rodents, with relatively low levels in the morning and high levels in the evening (Moore and Eichler, 1972). We assayed both AM and PM plasma CORT levels in the same mice taken two days before and two days after the CSDS/CR (**Fig. 2A).** We fit a linear mixed-effects model with CORT (ng/mL) levels as the outcome variable and pre- vs. post- CSDS/CR, time of day, and group and their interactions as the fixed effects. Across control, resilient, and susceptible mice, time of day was a significant predictor of CORT levels (*est*=1.552, *sd*=0.503, *p*=0.003), indicating that the diurnal rhythm of CORT remained robust in all groups. Importantly, this diurnal pattern was not predictive of social outcome (*p*>0.05), denoting that the resilient/susceptible phenotype did not shift the circadian cycle of stress hormones. Moreover, pre- and post-CSDS/CR levels were not predictive (*p*>0.05), suggesting that the process of going through the CSDS/CR itself did not alter baseline CORT levels (**Fig. 2D**).

### During the Initial Defeat Encounter, Susceptible Mice Engaged in More Escape Behavior

During agonistic encounters, mice engage in a variety of coping behaviors, including freeze, fight, and escape (McLaughlin et al., 2006; Scott, 1966). Therefore, we asked whether the type of coping behavior utilized during the defeat encounter has any predictive value for the resilient or susceptible outcome. To do this, we quantified these active and passive coping behaviors (**Table 1**) on Day 1 (a stress-naïve first encounter) and on Day 10 (experienced final stress encounter) (**Fig. 3A)**. As shown in **Fig. 3B**, resilient and susceptible mice displayed different patterns of freeze, fight, and escape. We then compared the expression of these behaviors on Day 1 and Day 10 between resilient and susceptible groups.

**Figure 3:**
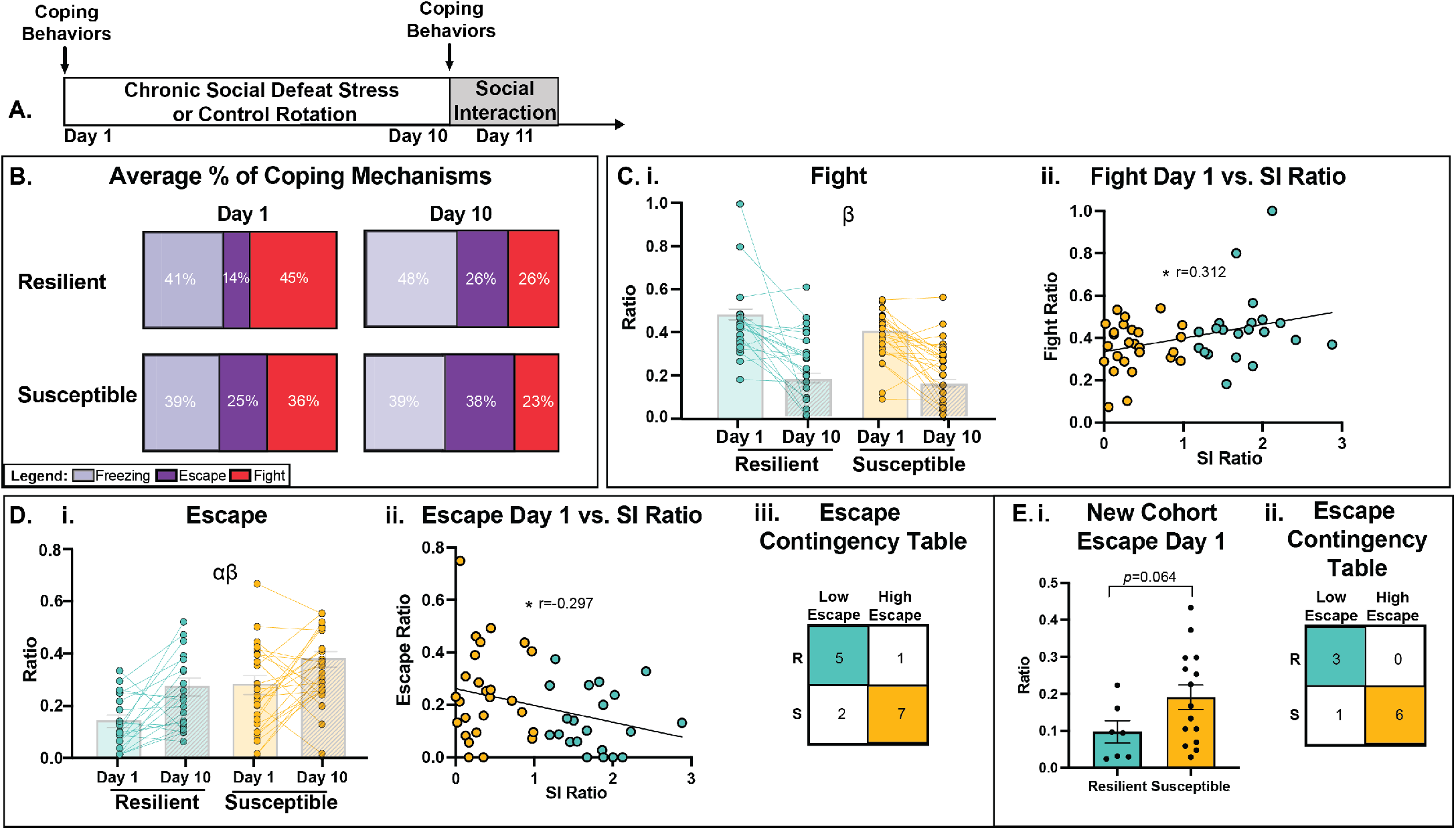
During Stress: Mice that end up susceptible used escape as a coping behavior. **3A.** Schematic timeline indicating observation of coping behaviors during Day 1 and Day 10 of defeat. **3B.** Average % of freezing, escape, and fight behaviors displayed on Day 1 and Day 10, according to resilient and susceptible phenotype. **3C. i**. Ratio of # of fights/total behaviors observed on Day 1 and Day 10 of defeat. Overall, fight behaviors decreased from Day 1 to Day 10, but levels of fighting were similar between resilient and susceptible groups. **3C. ii.** Fight ratios measured on Day 1 of defeat were positively correlated with SI ratios measured post-defeat. **3D. i.** Ratio of # of escapes/total behaviors observed on Day 1 and Day 10 of defeat. Escape behaviors were higher overall in susceptible compared to resilient mice and also higher overall on Day 10 compared to Day 1. **3D. ii.** Escape ratios measured on Day 1 of defeat were negatively correlated with SI ratios measured post-defeat. **3D. iii.** Escape contingency table illustrating a higher proportion of susceptible mice escape compared to resilient on Day 1. **3E. i.** Replication of Escape analysis in an additional cohort presented a trend for susceptible mice to escape more compared to resilient. **3E. ii.** Escape contingency table in replicate cohort illustrating a higher proportion of susceptible mice escape compared to resilient on Day 1. Significance Codes: Group Main Effect α, Day Main Effect β, \**p*<0.05. Colors: blue = resilient and yellow = susceptible.

#### Freezing behaviors were not related to resilient or susceptible outcomes

For all three freezing behaviors (Upright, Crouch, or Forward), there were no changes across Day or between Groups (**Supp. Fig. 1C-D, Supp. Table 5**; Day *p*>0.05, Group *p*>p0.05). Freezing behaviors were also not correlated with post-defeat SI ratios (**Supp. Table 15).** Thus, these data suggest that the freezing response is not related to the sociability outcome. Therefore, we assessed whether differences in the active coping behaviors (fight versus escape) mapped onto future expression of resilience and susceptibility.

#### Greater initial fighting was correlated with subsequent resilience

Fighting behavior over the course of the defeat protocol is dynamic. Over the course of the ten days of the defeat protocol, there was an overall significant decrease in fight duration (**Supp. Fig. 1B**; Test Day *F*1,44=16.20, *p*<0.001). Across all defeat sessions, neither fight duration nor ratio of total fight engagements differentiated the resilient and susceptible groups, as these groups displayed similar durations of active fighting across the defeat sessions (**Supp. Fig. 1B**; Group *F*_1,44_=0.030, *p*=0.860). Similarly, the fight ratios over the course of defeat were similar between resilient and susceptible mice (**Fig. 3Ci**; Group *F*_1,44_=3.45, *p*=0.070). However, the amount of fighting behavior displayed early in the course of defeat (in the initial encounter) was related to the eventual sociability phenotype, as indicated by a significant positive correlation between SI ratio and fight behavior on Day 1 (**Fig. 3Cii**; *r*=0.312, *p*=0.031). Thus, although overall fighting levels were similar between groups, higher SI ratios correlated with higher levels of fighting on the first aggressive encounter.

#### Day 1 escape behavior predicted which mice become susceptible

Escape behaviors are also dynamic, but in contrast to fight, escape ratios increased over the course of defeat (**Fig 3D. i.**; Test Day *F*_2,44_=29.92, *p*<0.001). There was also a significant group difference for this coping behavior, as mice that emerged as susceptible exhibited significantly greater escape behaviors on Day 1 than resilient mice (**Fig. 3D. i.**; Group *F*_1,44_=7.070, *p*=0.011). Notably, we observed a significant negative correlation between SI Ratio and escape behavior on Day 1 (**Fig. 3D. ii.**; *r*=-0.297, *p*=0.040).

We next asked whether escape behavior could predict susceptibility/resilience prospectively. To do this, we separated mice on Day 1 into two groups: high escape vs. low escape, based on whether mice escaped one standard deviation above or below the mean (*mea*n=0.195, *sd*=+/−0.160). A Fisher Exact Test revealed a significant correlation between SI Ratio and high/low escape behavior (*p*=0.041). A 2×2 contingency table analysis shows that on Day 1, the proportion of high escape mice that become susceptible to CSDS (7 out of 9) was larger than resilient individuals (1 out of 6) (**Fig. 3D. iii.)**.

We used an additional cohort to verify that Day 1 high escape behaviors predict the emergence of susceptibility. In this cohort, mice overall escaped at an average ratio of 0.161 (*sd*= +/− 0.117) with a trend for future susceptible mice to have a higher escape ratio (**Fig. 3E. i.**; *p*=0.064). After separating this cohort into high vs. low escape, a Fisher Exact Test revealed again that there was a significant correlation between SI Ratio and high/low escape behavior (*p*=0.033). A 2×2 contingency table analysis shows that on Day 1, the proportion of high escape mice that become susceptible to CSDS (6 out of 7) was larger than resilient individuals (0 out of 3) (**Fig. 3E. ii)**. Thus, examining escape behavior as either a continuous measure or categorical variable shows a predictive relationship between high escape and eventual susceptibility.

### Post-Stress: Comprehensive Behavioral Phenotyping Associated with Social Outcome

#### Traditional Affective Readouts: The Open Field (OF) and Forced Swim Test (FST)

To address whether resilient and susceptible phenotypes (according to the SI test) are generalizable to traditional measures of anxiety- and depressive-like behavior, we conducted the Open Field (OF, anxiety-like measure) and Forced Swim Test (FST, depression-like measure) following CSDS or CR (**Fig. 4A**).

**Figure 4:**
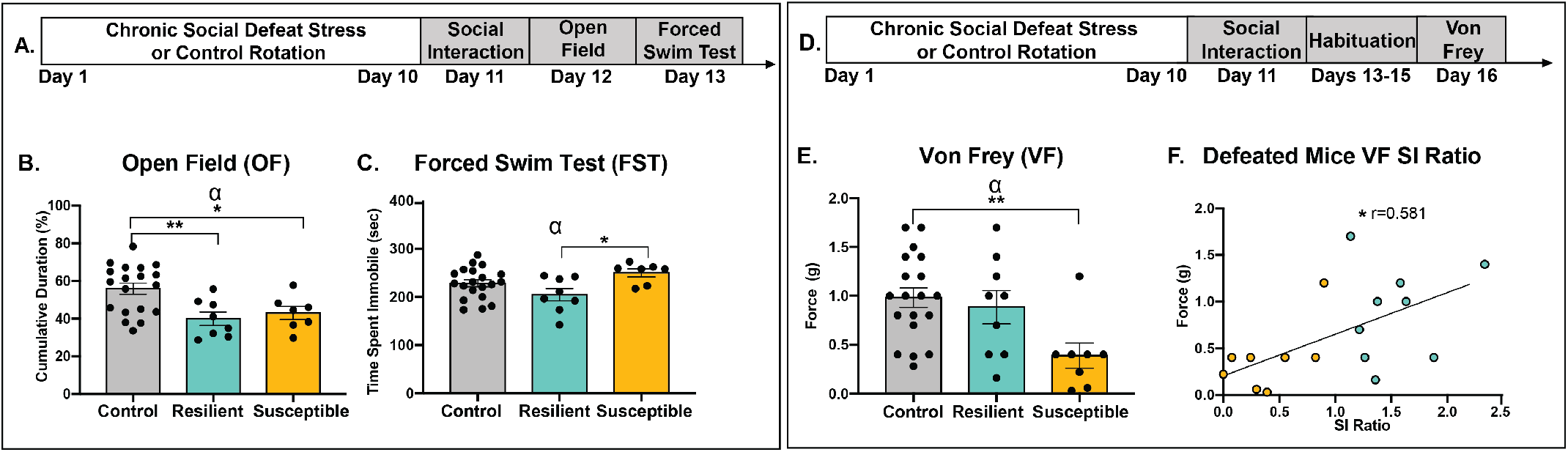
Post-Stress: Characterization of Behavioral Outcomes. **4A**. Schematic timeline of assessing traditional read-outs of anxiety-like (Open field, OF) and depression-like (Forced swim test (FST) behavior tests. **4B**. Cumulative duration (%) of time spent in the open area of the OF was overall decreased by defeat. **4C.** Resilient mice spent less time immobile compared to susceptible mice. **4D.** Schematic timeline of Von Frey (VF) testing. **4E.** In the VF test, susceptible mice had increased pain sensitivity compared to controls. **4F**. Lower SI ratios were correlated with lower force (g) filaments used to elicit a response in the VF test (increased pain sensitivity). Significance Codes: Group Main Effect α, \**p*<0.05, \*\**p*< 0.01. Colors: grey = control, blue = resilient, yellow = susceptible.

In the OF test, social defeat reduced the percent time spent in center compared to controls (**Fig. 4B**; *F*_2,31_=6.689, *p*=0.004), indicating a general increase in anxiety-like behavior following CSDS. However, there was no difference between resilient and susceptible mice (*p*>0.05), and there was no correlation between SI Ratios and percent time spent in the center in defeated mice (*p*>0.05) (**Supp. Table 10**). Therefore, although defeat itself increased anxiety-like behavior in the OF, there was no difference between resilient and susceptible groups.

In the FST, there was a significant difference across groups in time spent immobile (**Fig. 4C**; *F*_2,31_=3.93, *p*=0.030). Compared to controls, there was no significant difference between resilient (*p>*0.05) or susceptible (*p>*0.05) mice. However, susceptible mice spent significantly more time immobile compared to resilient mice (*p*=0.023). In this test, SI Ratios did not correlate with immobility time in defeated mice (**Supp. Table 10**; *p*>0.05).

#### Mice with lower SI Ratios displayed greater sensitivity to mechanical nociception **(Fig. 4D)**

Due to the physical nature of social defeat, we conducted the Von Frey (VF) test to understand whether pain sensitivity differed between resilient and susceptible mice. There was a significant effect of group on filament force (g) used in the VF test (**Fig. 4E**; *F*_2,33_=5.163, *p*=0.011). Susceptible mice needed a significantly lower filament force to elicit a mechanical response than controls, indicating an increased mechanical nociception sensitivity (*p*=0.009). In contrast, resilient mice had a similar filament force threshold to control mice (*p*>0.05). There was a trend for lower pain threshold in susceptible mice compared to resilient (*p*=0.069). And moreover, within defeated mice, there was a positive correlation between SI Ratio and force (g) of filament threshold (**Fig. 4F**; *p*=0.015, *r*=0.581), indicating a relationship between social avoidance and increased pain sensitivity. These data highlight a relationship between social interaction and pain sensitivity that goes beyond the traditional measures of affective-like behaviors.

### Post-Stress: Sociability Outcome was Related to Physiological Measures of Weight and CORT

To determine whether there were neuroendocrine and physiological differences in resilient and susceptible mice following stress, we conducted a social stress challenge and observed changes in body weight and CORT.

#### In response to a social stress challenge, lower SI ratios were correlated with lower CORT expression

Although pre-defeat experiments showed no differences in basal CORT rhythms, we wanted to determine whether resilient and susceptible mice would display different CORT responses when given a social stress challenge. Following 40-minute exposure to a CD1 mouse, there was an overall group difference between control, resilient, and susceptible mice (**Fig. 5B**; *F*_2,33_=4.560, *p*=0.0214). This group effect was driven by increased CORT expression in resilient mice compared to controls (*p*=0.016). Interestingly, within defeated mice, there was a significant positive correlation between SI ratios and CORT levels (*r*=0.055, *p*=0.026), indicating that the more socially interactive mice mounted higher CORT responses to the social stress challenge (**Fig. 5C**).

**Figure 5:**
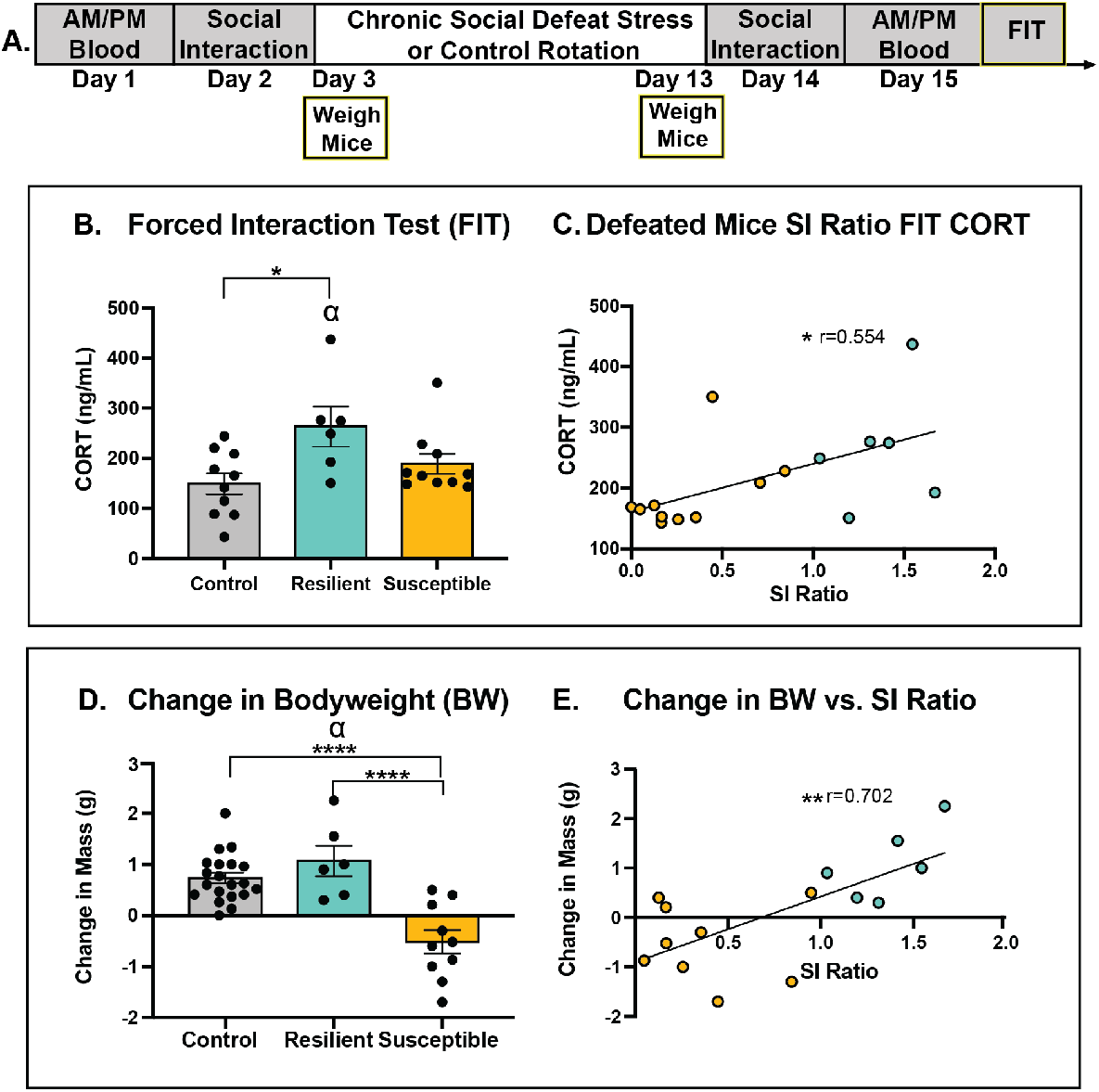
Susceptible mice gained less weight and mounted a blunted CORT response compared to resilient mice. **5A**. Schematic timeline of body weight and Forced Interaction Test (FIT) CORT measurements. **5B.** FIT-induced CORT (ng/mL) levels increased in resilient mice compared to controls. **5C.** In defeated mice, lower FIT CORT (ng/mL) levels correlated with lower SI Ratios. **5D.** Change in BW (Day 10 – Day 1) indicates that susceptible mice had reduced weight gain compared to resilient and control mice. **5E.** In defeated mice, higher SI ratios were significantly correlated with larger weight gains over the course of the experiment. Significance Codes: Group Main Effect α, *p<0.05, **p< 0.01. Colors: grey = control, blue = resilient, yellow = susceptible.

#### Susceptible mice had reduced weight gain by the end of CSDS

As an additional physiological measure of the effects of stress, we compared changes in body weight across the experimental period in mice exposed to CSDS or CR (**Fig. 5A**). Weight change across social defeat was calculated (Day 10 weight – Day 1 weight) and showed an overall group difference between control, resilient, and susceptible mice (**Fig. 5D**; *F*_2,33_=18.60, *p*<0.001). This difference was driven by a significant reduction in weight in susceptible mice compared to both control (*p*<0.001) and resilient (*p*<0.001) mice. Within the defeated group, there was a significant positive correlation between SI Ratio and body weight (**Fig. 5E**; *r*=0.492, *p*<0.001), further indicating that social avoidance was related to reduced weight gain.

We ran a linear regression to test whether there was a relationship between body weight and SI Ratio on FIT CORT expression. We found that while there was an effect of body weight change on SI Ratio (*p*=0.017), there was no effect of weight change on FIT CORT expression (**Supp. Table 9;** *p*>0.05). This analysis indicates that although body weight and stress-induced CORT were independently related to social reactivity outcome of defeat, the two physiological variables were not directly related to each other.

### Post-Stress: Susceptible Mice Exhibited an Ability to Discriminate Social Threat

An important remaining question about the CSDS model was whether the reduced SI ratios observed in susceptible mice reflect a generalized reduction in social motivation or whether the change is specific to the strain associated with defeat. To determine the specificity of social interaction behavior to the aggressor strain (CD1), two additional cohorts of defeated and control mice were assessed with SI tests using different strains of mice as social partners. First, to measure social responses to animals *not* associated with an aggressive encounter, subjects were tested with either a novel C57BL/6J mouse or a mouse from a different outbred strain (Black Swiss). These “non-aggressor” tests were followed by a SI test using a CD1 mouse as a social partner. (**Fig. 6A)**.

**Figure 6:**
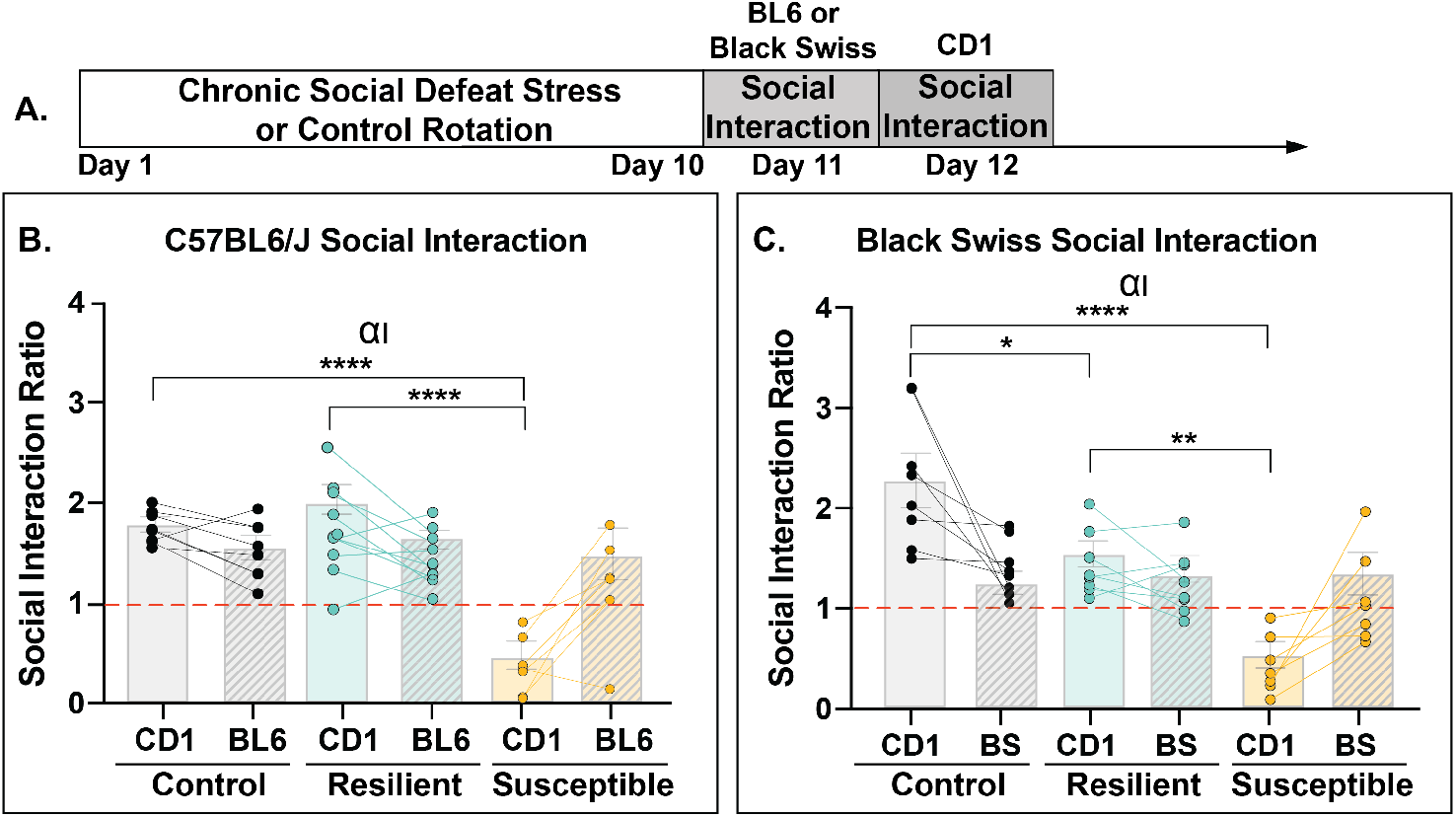
Susceptibility in the SI test was specific to strain used during defeat. **6A.** Schematic timeline of strain-specificity social interaction testing. **6B.** When tested with the aggressor strain (CD1) as the social partner, susceptible mice had reduced SI ratios compared to the other groups, indicating social avoidance. However, susceptible mice displayed similar SI ratios compared to controls and resilient mice when placed with a C56BL/6J (BL6). **6C.** When tested with a Black Swiss (BS), susceptible mice displayed SI ratios similar to control and resilient mice. Susceptible mice only exhibited social avoidance when a CD1 was the social partner in the SI test. Resilient mice displayed lower SI ratios compared to controls in the CD1 interaction but remained above 1 (socially interactive). Significance Codes: Group Main Effect α, Interaction Effect I, *p<0.05, **p< 0.01, p<****0.0001. Colors: red dash= SI ratio of 1, grey = control, blue = resilient, yellow = susceptible.

#### Susceptible mice were not socially avoidant when tested with non-aggressor strains

When we tested defeat subjects with a partner of the same strain (C57BL/6J) or with a CD1, we found that there was a significant interaction effect between group and strain of social partner (**Fig. 6B**; *F*_2,23_=13.70, *p*<0.001). This interaction reflects a group difference in the CD1 SI test, but not the C57BL/6J SI test (CD1 test: *F*_2,23_=34.08, *p*<0.001; C57BL/6J test: *F*_2,23_=0.373, *p*=0.693). When the CD1 was the social partner, susceptible mice displayed significantly lower SI Ratios compared to both resilient (*p*<0.001) and control (**Supp. Table 13**; *p*<0.001). In contrast, when the C57BL/6J was the social partner, susceptible, resilient, and control mice all had comparable SI Ratios (**Supp. Table 13**; all post-hoc comparisons: *p>*0.05).

When we tested subjects with a different outbred strain (Black Swiss) or with a CD1, we again found a significant interaction between group and strain of social partner (**Fig. 6B**; *F*_2,20_=13.27, *p*<0.001). Similarly, this interaction reflects a group difference in the CD1 SI test, but not the Black Swiss SI test (CD1 test: *F*_2,20_=19.39, *p*<0.001; Black Swiss test: *F*_2,20_=0.120, *p*=0.888). When the CD1 was the social partner, susceptible mice displayed significantly lower SI Ratios compared to both resilient (*p*<0.001) and control (*p*<0.001) mice. Resilient mice displayed a significant decrease in SI ratios compared to controls (*p*=0.035); nevertheless, resilient mice tested with a CD1 still had SI ratios above 1, which is indicative of a socially interactive phenotype (one-sample *t*-test, *t*=3.801, df=7, *p*=0.007). Importantly, when the Black Swiss was the social partner in the SI test, susceptible, resilient, and control mice all had comparable SI Ratios (**Supp. Table 13**; all post-hoc comparisons: *p>*0.05).

Collectively, our results demonstrate that, when tested with either of the non-CD1 (non-aggressor) strains, susceptible mice exhibited SI ratios comparable to control and resilient mice, indicating that the shift in sociability is specific to strains associated with the actual defeat. Therefore, susceptible mice can clearly discriminate between a threatening strain (CD1) and non-threatening (C57BL6/J or Black Swiss) strain.

## DISCUSSION

The CSDS model has proven valuable in investigating the underlying neurobiological mechanisms of social reactivity after resilient and susceptible groups are established. Following CSDS, neuroanatomical differences (Anacker et al., 2016), transcriptional profiles (Bagot et al., 2016), and neural activity patterns (Hultman et al., 2018; Muir et al., 2020, 2018) have all been linked to susceptibility or resilience. Our work adds a characterization of the behavioral differences either before or early in the stress experience that may set the course for the social outcome. We assessed behavioral, neuroendocrine, and physiological differences prestress, during the early social stress experience, and post-stress. Collectively, the current work outlines several key findings that shed light on the affective makeup of resilient and susceptible mice.

### Key Findings

a. We did not identify a priori behavioral readouts that predicted resilient or susceptible outcomes. The degree of social interaction before stress is not predictive of social response following stress, suggesting that these animals do not simply differ in “sociability.” Moreover, physiologically, basal CORT has no predictive value for social reactivity to CSDS.
b. By contrast, *coping style during the first encounter* is a key variable in the emergence of the eventual phenotype: higher rates of escape behavior on Day 1 indicate a greater likelihood of a subsequent classification of susceptible. In addition, the propensity for more fighting on Day 1 correlates with greater resilience scores.
c. Following stress, social avoidance is not generalizable to traditional measures of affective behavior. However, it is associated with greater sensitivity to physical pain and significant weight loss. Moreover, lower social interaction ratios correlate with lower CORT expression when mice are placed back in the social stress context.
d. Social avoidance is social context-specific, as susceptible mice do not avoid non-aggressive strains. This finding suggests that threat discrimination may play a key role in social reactivity following CSDS.

Taken together, our findings demonstrate that the susceptible, socially avoidant animals exhibit high reactivity to a specific social stressor, characterized by a greater propensity to escape during the early encounters. This initial difference evolves into a relatively lower endocrine stress response to social threat, along with greater sensitivity to pain and weight loss. However, susceptible mice appear clearly attuned to the nature of the aggressor and do not generalize the threat response to either con-specifics or other novel strains of mice. Through this work, we have expanded our understanding of susceptibility in this animal model, adding context to the socially avoidant phenotype.

#### Evaluating how changes in social motivation track with traditional measures of emotional vulnerability

Across multiple labs, the variation of behavior in response to social defeat reflects the complexity of defining features of vulnerability. When assessed after the defeat, traditional measures of anxiety-like behavior, such as open field and elevated plus maze, generally do not track with resilient and susceptible outcomes (Krishnan et al., 2007). There have been contradictory findings related to the existence of group differences in the forced swim test (Krishnan et al., 2007; Lee et al., 2021; Zhang et al., 2021). Although we detected a difference in immobility between resilient and susceptible mice, both groups were comparable to controls. Additionally, within defeated animals, there was no relationship between social reactivity and this particular depressive-like measure. It is important to highlight that CSDS is both a physical and psychosocial stressor—in our work, delineation of resilient and susceptible animals involved measures that were physical (pain sensitivity and body weight) or psychosocial (coping behavior, FIT CORT, and susceptible threat-discrimination). Thus, although anxiety- or depressive-like assays can be useful in characterizing general affective states following CSDS, these tests do not appear to reflect the relevant stimuli being encoded by the social stress itself and may be less tied to the social outcome.

#### Predicting the resilient-susceptible outcome

Recently there has been greater emphasis on uncovering what traits or combinations of traits are needed to predict resilient and susceptible outcomes. For example, Nasca *et al.* found that predicting susceptibility is optimized when multiple behavioral, neuroanatomical, and physiological measures (Nasca et al., 2019). When looking specifically at social behavior features, Larrieu *et al.* identified that hierarchal status before defeat predicts the social phenotype; mice that are higher-ranked in their home cage are more likely to exhibit susceptible and socially avoidant behaviors following social defeat (Larrieu et al., 2017). Additionally, Milic et al. identified that low exploratory behavior and heightened fear-learning in a foot-shock paradigm could predict susceptibility (Milic et al., 2021). The identification of various predictive traits of social outcome reflects the overall complexity of individual variation in response to social stress. Our detailed social profiling contributes an additional insight, as we specifically focused on asking whether there was a feature of the defeat itself that could predict the final social outcome. We provide novel evidence that, on Day 1 of social defeat, we can predict which mice will be subsequently classified as susceptible or resilient based on high or low rates of escape behavior. Specifically, mice that escaped at significantly higher rates were more likely to develop susceptibility. These data show for the first time that pre-existing differences in coping strategies may represent a useful predictive factor for whether resiliency or susceptibility develops.

#### Evaluating the cost-benefit balance in susceptible mice

The two main coping behaviors we observed during the defeat, fight and escape, can be understood in the context of evolutionarily conserved predator-prey interactions (Koolhaas et al., 1999). For example, mice can either engage in the fight to assert active territorial control (Anderson and Hill, 1965) or deploy an escape strategy to avoid the looming threat of the approaching predator (De Franceschi et al., 2016; Eilam, 2005; Shang et al., 2018). Future resilient mice appear proactive and may be attempting to assert dominance over the social threat through heightened aggression and fight engagements. This interpretation is further supported by the finding that CORT responses to social stress challenge were elevated in resilient mice, as increased plasma glucocorticoids have been linked to increased levels of aggression and fight engagements (Haller et al., 1998). Notably, this heightened level of aggression may come at a cost, as sustained CORT expression can have detrimental health effects on both body and brain (McEwen and Akil, 2020). Future studies analyzing the temporal nature of CORT responses in resilient vs. susceptible mice are needed to evaluate the nature of stress-induced changes to the function of the HPA axis.

Although susceptibility has often been thought to reflect a maladaptive reduction in social motivation, our results suggest possible advantages to this response. In susceptible mice, the high propensity to escape indicates that these animals tend to employ an evasive strategy to avoid harm and are perhaps highly sensitive to threat. Interestingly, we found susceptible mice do not appear to have reduced social motivation in general, only reduced exploration specific to the strain associated with aggression. Similar to our findings, when using a modified social interaction test in which defeated mice were placed in a three-chambered social approach task, Ayash *et al.* found that susceptible mice interacted significantly less with a novel CD-1 than a C57BL6/J or 129/Sv (Ayash et al., 2020). The potential benefits of susceptibility, in terms of heightened threat discrimination, appear to come at some cost, as these animals also had reduced weight gain and increased pain sensitivity following CSDS. Other studies have observed overall increased pain sensitivity following defeat (Marco Pagliusi et al., 2020; M Pagliusi et al., 2020), but it is unclear whether individual variation in pain sensitivity precedes the defeat; additional studies are needed to uncover whether this is a predictive factor. Thus, as defined in the context of CSDS, susceptibility should be viewed in a nuanced way, in terms of costbenefit, based on the conditions in which the behavior is expressed. As such, the ethological basis of susceptibility may reflect underlying differences in threat assessment and discrimination tactics.

#### Concluding Remarks

It is worth recalling that resilient and susceptible mice in these and other studies derive from a single inbred strain and therefore share a common genetic makeup. Despite this, inbred strains exhibit a high degree of individual variability in behavioral outputs (Tuttle et al., 2018; Wahlsten et al., 2006), highlighting the key role of developmental and contextual variables in shaping emotional reactivity. Some of these variables may arise from maternal behavior (Pedersen et al., 2011) or from the social context in the animal cage leading to social dominance hierarchies (Horii et al., 2017). Here, our longitudinal study suggests the existence of an ongoing, dynamic interplay between the animal’s initial propensity for adopting different coping strategies, the specific characteristics of the stress condition, and the cumulative effects of daily stress experience. Over the course of the stress, these factors work together to shape the outcome of becoming resilient or susceptible, each with their own set of costs and benefits.

This perspective provides a context for future studies investigating the neural correlates associated with the early differences in coping strategies, as well as the unique dynamic changes related to the emergence of distinct social stress phenotypes. Uncovering the neural mechanisms associated with coping during social stress can enhance our understanding of human affective and stress disorders to inform treatment and prevention strategies.

## REFERENCES

Abdallah, C.G., Geha, P., 2017. Chronic pain and chronic stress: two sides of the same coin? Chronic Stress (Thousand Oaks) 1. doi:10.1177/2470547017704763

Adam, E.K., Quinn, M.E., Tavernier, R., McQuillan, M.T., Dahlke, K.A., Gilbert, K.E., 2017. Diurnal cortisol slopes and mental and physical health outcomes: A systematic review and meta-analysis. Psychoneuroendocrinology 83, 25–41. doi:10.1016/j.psyneuen.2017.05.018

Akil, H., 2005. Stressed and depressed. Nat. Med. 11, 116–118. doi:10.1038/nm0205-116

Anacker, C., Scholz, J., O’Donnell, K.J., Allemang-Grand, R., Diorio, J., Bagot, R.C., Nestler, E.J., Hen, R., Lerch, J.P., Meaney, M.J., 2016. Neuroanatomic differences associated with stress susceptibility and resilience. Biol. Psychiatry 79, 840–849. doi:10.1016/j.biopsych.2015.08.009

Anderson, P.K., Hill, J.L., 1965. Mus musculus: Experimental Induction of Territory Formation. Science 148, 1753–1755. doi:10.1126/science.148.3678.1753

Ayash, S., Schmitt, U., Müller, M.B., 2020. Chronic social defeat-induced social avoidance as a proxy of stress resilience in mice involves conditioned learning. J. Psychiatr. Res. 120, 64–71. doi:10.1016/j.jpsychires.2019.10.001

Bagot, R.C., Cates, H.M., Purushothaman, I., Lorsch, Z.S., Walker, D.M., Wang, J., Huang, X., Schlüter, O.M., Maze, I., Peña, C.J., Heller, E.A., Issler, O., Wang, M., Song, W.-M., Stein, J.L., Liu, X., Doyle, M.A., Scobie, K.N., Sun, H.S., Neve, R.L., Geschwind, D., Dong, Y., Shen, L., Zhang, B., Nestler, E.J., 2016. Circuit-wide Transcriptional Profiling Reveals Brain Region-Specific Gene Networks Regulating Depression Susceptibility. Neuron 90, 969–983. doi:10.1016/j.neuron.2016.04.015

Billings, A.G., Moos, R.H., 1984. Coping, stress, and social resources among adults with unipolar depression. J. Pers. Soc. Psychol. 46, 877–891. doi:10.1037//0022-3514.46.4.877

Butler, R.K., Finn, D.P., 2009. Stress-induced analgesia. Prog. Neurobiol. 88, 184–202. doi:10.1016/j.pneurobio.2009.04.003

Connor-Smith, J.K., Compas, B.E., 2002. Vulnerability to Social Stress: Coping as a Mediator or Moderator of Sociotropy and Symptoms of Anxiety and Depression. Springer Science and Business Media LLC. doi:10.1023/a:1013889504101

De Franceschi, G., Vivattanasarn, T., Saleem, A.B., Solomon, S.G., 2016. Vision guides selection of freeze or flight defense strategies in mice. Curr. Biol. 26, 2150–2154. doi:10.1016/j.cub.2016.06.006

Eilam, D., 2005. Die hard: a blend of freezing and fleeing as a dynamic defense--implications for the control of defensive behavior. Neurosci. Biobehav. Rev. 29, 1181–1191. doi:10.1016/j.neubiorev.2005.03.027

Fox, J., Weisberg, S., 2019. An {R} Companion to Applied Regression, Third Edition [WWW Document]. URL https://socialsciences.mcmaster.ca/jfox/Books/Companion/

Golden, S.A., Covington, H.E., Berton, O., Russo, S.J., 2011. A standardized protocol for repeated social defeat stress in mice. Nat. Protoc. 6, 1183–1191. doi:10.1038/nprot.2011.361

Grosjean, P., Ibanez, F., 2018. pastecs: Package for Analysis of Space-Time Ecological Series [R package version 1.3.21] [WWW Document]. URL https://cran.r-project.org/web/packages/pastecs/index.html

Haller, J., Halasz, J., Makara, G.B., Kruk, M.R., 1998. Acute effects of glucocorticoids: behavioral and pharmacological perspectives. Neurosci. Biobehav. Rev. 23, 337–344. doi:10.1016/s0149-7634(98)00035-9

Hollis, F., Kabbaj, M., 2014. Social defeat as an animal model for depression. ILAR J. 55, 221–232. doi:10.1093/ilar/ilu002

Horii, Y., Nagasawa, T., Sakakibara, H., Takahashi, A., Tanave, A., Matsumoto, Y., Nagayama, H., Yoshimi, K., Yasuda, M.T., Shimoi, K., Koide, T., 2017. Hierarchy in the home cage affects behaviour and gene expression in group-housed C57BL/6 male mice. Sci. Rep. 7, 6991. doi:10.1038/s41598-017-07233-5

Hultman, R., Ulrich, K., Sachs, B.D., Blount, C., Carlson, D.E., Ndubuizu, N., Bagot, R.C., Parise, E.M., Vu, M.-A.T., Gallagher, N.M., Wang, J., Silva, A.J., Deisseroth, K., Mague, S.D., Caron, M.G., Nestler, E.J., Carin, L., Dzirasa, K., 2018. Brain-wide Electrical Spatiotemporal Dynamics Encode Depression Vulnerability. Cell 173, 166–180.e14. doi:10.1016/j.cell.2018.02.012

Iio, W., Takagi, H., Ogawa, Y., Tsukahara, T., Chohnan, S., Toyoda, A., 2014. Effects of chronic social defeat stress on peripheral leptin and its hypothalamic actions. BMC Neurosci. 15, 72. doi:10.1186/1471-2202-15-72

Jeong, J.Y., Lee, D.H., Kang, S.S., 2013. Effects of chronic restraint stress on body weight, food intake, and hypothalamic gene expressions in mice. Endocrinol Metab (Seoul) 28, 288–296. doi:10.3803/EnM.2013.28.4.288

Koolhaas, J.M., Korte, S.M., De Boer, S.F., Van Der Vegt, B.J., Van Reenen, C.G., Hopster, H., De Jong, I.C., Ruis, M.A., Blokhuis, H.J., 1999. Coping styles in animals: current status in behavior and stress-physiology. Neurosci. Biobehav. Rev. 23, 925–935. doi:10.1016/s0149-7634(99)00026-3

Krishnan, V., Han, M.-H., Graham, D.L., Berton, O., Renthal, W., Russo, S.J., Laplant, Q., Graham, A., Lutter, M., Lagace, D.C., Ghose, S., Reister, R., Tannous, P., Green, T.A., Neve, R.L., Chakravarty, S., Kumar, A., Eisch, A.J., Self, D.W., Lee, F.S., Tamminga, C.A., Cooper, D.C., Gershenfeld, H.K., Nestler, E.J., 2007. Molecular adaptations underlying susceptibility and resistance to social defeat in brain reward regions. Cell 131, 391–404. doi:10.1016/j.cell.2007.09.018

Larrieu, T., Cherix, A., Duque, A., Rodrigues, J., Lei, H., Gruetter, R., Sandi, C., 2017. Hierarchical status predicts behavioral vulnerability and nucleus accumbens metabolic profile following chronic social defeat stress. Curr. Biol. 27, 2202–2210.e4. doi:10.1016/j.cub.2017.06.027

Lee, C.-W., Fang, Y.-P., Chu, M.-C., Chung, Y.-J., Chi, H., Tang, C.-W., So, E.C., Lin, Hsin-Chuan, Lin, Hui-Ching, 2021. Differential mechanisms of synaptic plasticity for susceptibility and resilience to chronic social defeat stress in male mice. Biochem. Biophys. Res. Commun. 562, 112–118. doi:10.1016/j.bbrc.2021.05.064

Lenth, R., 2020. Estimated Marginal Means, aka Least-Squares Means. R package version 1.5.0 [WWW Document]. URL https://CRAN.R-project.org/package=emmeans

McEwen, B.S., Akil, H., 2020. Revisiting the stress concept: implications for affective disorders. J. Neurosci. 40, 12–21. doi:10.1523/JNEUROSCI.0733-19.2019

McLaughlin, J.P., Li, S., Valdez, J., Chavkin, T.A., Chavkin, C., 2006. Social defeat stress-induced behavioral responses are mediated by the endogenous kappa opioid system. Neuropsychopharmacology 31, 1241–1248. doi:10.1038/sj.npp.1300872

Milic, M., Schmitt, U., Lutz, B., Müller, M.B., 2021. Individual baseline behavioral traits predict the resilience phenotype after chronic social defeat. Neurobiol. Stress 14, 100290. doi:10.1016/j.ynstr.2020.100290

Moore, R.Y., Eichler, V.B., 1972. Loss of a circadian adrenal corticosterone rhythm following suprachiasmatic lesions in the rat. Brain Res. 42, 201–206. doi:10.1016/0006-8993(72)90054-6

Muir, J., Lorsch, Z.S., Ramakrishnan, C., Deisseroth, K., Nestler, E.J., Calipari, E.S., Bagot, R.C., 2018. In vivo fiber photometry reveals signature of future stress susceptibility in nucleus accumbens. Neuropsychopharmacology 43, 255–263. doi:10.1038/npp.2017.122

Muir, J., Tse, Y.C., Iyer, E.S., Biris, J., Cvetkovska, V., Lopez, J., Bagot, R.C., 2020. Ventral Hippocampal Afferents to Nucleus Accumbens Encode Both Latent Vulnerability and Stress-Induced Susceptibility. Biol. Psychiatry. doi:10.1016/j.biopsych.2020.05.021

Nasca, C., Menard, C., Hodes, G., Bigio, B., Pena, C., Lorsch, Z., Zelli, D., Ferris, A., Kana, V., Purushothaman, I., Dobbin, J., Nassim, M., DeAngelis, P., Merad, M., Rasgon, N., Meaney, M., Nestler, E.J., McEwen, B.S., Russo, S.J., 2019. Multidimensional predictors of susceptibility and resilience to social defeat stress. Biol. Psychiatry 86, 483–491. doi:10.1016/j.biopsych.2019.06.030

Pagliusi, M, Bonet, I.J.M., Brandão, A.F., Magalhães, S.F., Tambeli, C.H., Parada, C.A., Sartori, C.R., 2020. Therapeutic and Preventive Effect of Voluntary Running Wheel Exercise on Social Defeat Stress (SDS)-induced Depressive-like Behavior and Chronic Pain in Mice. Neuroscience 428, 165–177. doi:10.1016/j.neuroscience.2019.12.037

Pagliusi, Marco, Bonet, I.J.M., Lemes, J.B.P., Oliveira, A.L.L., Carvalho, N.S., Tambeli, C.H., Parada, C.A., Sartori, C.R., 2020. Social defeat stress-induced hyperalgesia is mediated by nav 1.8+ nociceptive fibers. Neurosci. Lett. 729, 135006. doi:10.1016/j.neulet.2020.135006

Pedersen, C.A., Vadlamudi, S., Boccia, M.L., Moy, S.S., 2011. Variations in Maternal Behavior in C57BL/6J Mice: Behavioral Comparisons between Adult Offspring of High and Low Pup-Licking Mothers. Front. Psychiatry 2, 42. doi:10.3389/fpsyt.2011.00042

Pinhero, J., Bates, D., S, D., D, S., R Core Team, 2020. nlme: Linear and Nonlinear Mixed Effects Models [R package version 3.1-153] [WWW Document]. URL https://cran.r-project.org/web/packages/nlme/index.html

Scott, J.P., 1966. Agonistic behavior of mice and rats: a review. Am Zool 6, 683–701. doi:10.1093/icb/6.4.683

Shang, C., Chen, Z., Liu, A., Li, Y., Zhang, J., Qu, B., Yan, F., Zhang, Y., Liu, W., Liu, Z., Guo, X., Li, D., Wang, Y., Cao, P., 2018. Divergent midbrain circuits orchestrate escape and freezing responses to looming stimuli in mice. Nat. Commun. 9, 1232. doi:10.1038/s41467-018-03580-7

Singmann, H., Bolker, B., Westfall, J., Aust, F., 2016. afex: Analysis of Factorial Experiments [R package version 1.0-1] [WWW Document]. afex: Analysis of Factorial Experiments. R package version 0.16-1. URL https://cran.r-project.org/web/packages/afex/index.html

Tuttle, A.H., Philip, V.M., Chesler, E.J., Mogil, J.S., 2018. Comparing phenotypic variation between inbred and outbred mice. Nat. Methods 15, 994–996. doi:10.1038/s41592-018-0224-7

Wahlsten, D., Bachmanov, A., Finn, D.A., Crabbe, J.C., 2006. Stability of inbred mouse strain differences in behavior and brain size between laboratories and across decades. Proc. Natl. Acad. Sci. USA 103, 16364–16369. doi:10.1073/pnas.0605342103

Wood, S.K., Bhatnagar, S., 2015. Resilience to the effects of social stress: evidence from clinical and preclinical studies on the role of coping strategies. Neurobiol. Stress 1, 164–173. doi:10.1016/j.ynstr.2014.11.002

Young, E.A., Lopez, J.F., Murphy-Weinberg, V., Watson, S.J., Akil, H., 2000. Hormonal evidence for altered responsiveness to social stress in major depression. Neuropsychopharmacology 23, 411–418. doi:10.1016/S0893-133X(00)00129-9

Zhang, K., Sakamoto, A., Chang, L., Qu, Y., Wang, S., Pu, Y., Tan, Y., Wang, X., Fujita, Y., Ishima, T., Hatano, M., Hashimoto, K., 2021. Splenic NKG2D confers resilience versus susceptibility in mice after chronic social defeat stress: beneficial effects of (R)-ketamine. Eur Arch Psychiatry Clin Neurosci 271, 447–456. doi:10.1007/s00406-019-01092-z

