## Supplemental Material for "Characterizing the Behavioral and Neuroendocrine Features of Susceptibility and Resilience to Social Stress"

### Supplemental Figures

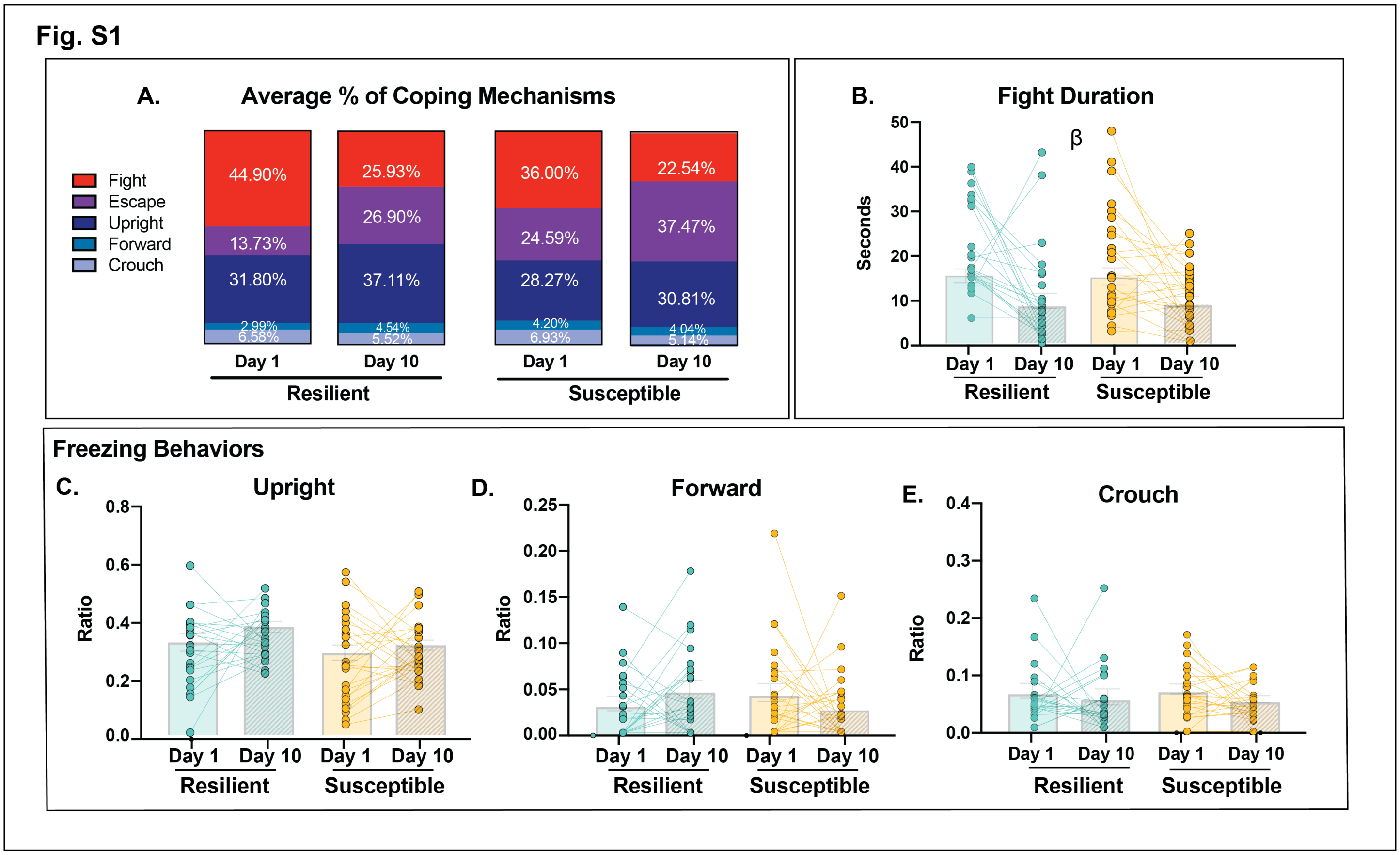


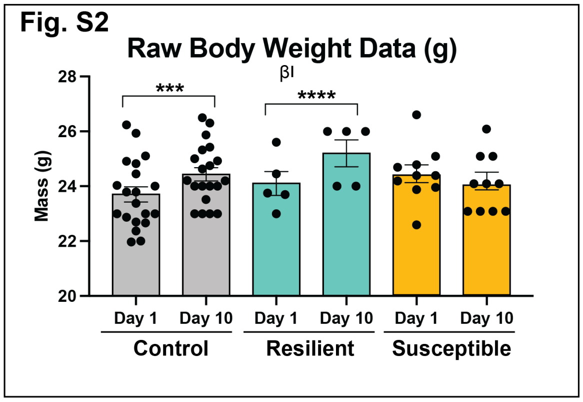


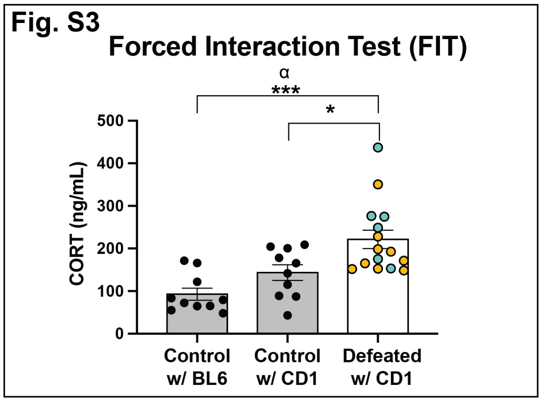


**Supplemental Figures**

**Figure S1: Coping Behaviors Used During Stress**

**S1A**.Parts of a whole bar chart representing the average percent of all coping behaviors used during defeat.

**S1B.** From Day 1 to Day 10, fight duration decreased similarly between resilient and susceptible groups.

**S1C**. Ratio of # of Upright Freezes/Total Behaviors Observed was similar between resilient and susceptible mice and did not change due to day.

**S1D.** Ratio of # of Forward Freezes/Total Behaviors Observed was similar between resilient and susceptible and did not change due to day.

**S1E**. Ratio of # of Crouch Back Freezes/Total Behaviors Observed was similar between resilient and susceptible and did not change due to day.

**Figure S2: Raw body weight data from Day 1 and Day 10 between control, resilient, and susceptible mice**.
There was a main effect of day and an interaction effect between group and day, as indicated by the significant increase in weight (g) in control and resilient mice, but not susceptible. Importantly, initial body weights were similar across groups (measured at the beginning of defeat).

**Figures S3: Forced Interaction Test.**
Mice were placed in similar conditions to the CR, CSDS, and SI test to look at CORT (ng/mL) expression. There was an overall Group main effect. Post-hoc Tukey tests revealed that CORT (ng/mL) expression of control mice across from BL6 or CD1 did not differ, but there was a significant increase in defeated mice CORT(ng/mL) expression compared to both control conditions.
Significance Codes: Group Main Effect α, Day Main Effect β, Interaction Effect ι, p<0.05, **p< 0.01, ***p< 0.001, p<****0.0001

### FIGURE 2 STATISTICS:

#### Table S1: Paired T-test Statistics within Group for Social Interaction Pre-Post Social Interaction

| Group Comparison | T(df) | P.Value |
|---|---|---|
| **Control Pre-Post** | **19(2.653)** | **0.0157** |
| **Defeat Pre-Post** | **15(3.855)** | **0.0016** |

#### Table S2: Mann-Whitney T-test Between Groups for Pre-Post Social Interaction

| Comparison | P.Value | Mann-Whitney U |
|---|---|---|
| Pre CR – Pre Defeat | P=0.2234 | 121 |
| **Post CR – Post Defeat** | **P=0.0035** | **70** |

#### Table S3: Pre-Post Pearson Correlations Table

cor.test(dataset$SIRatioPre, dataset$SIRatioPost, method=Pearson)

| Behavior | Comparison | Pearson r | P.Value |
|---|---|---|---|
| **Pre-Post Social Interaction Test** | | | |
| | Control Rotation Pre vs. Post | 0.1900 | 0.3988 |
| | Defeat Pre vs. Post | -0.3616 | 0.1688 |

#### Table S4: Baseline CORT differences

1) We looked at the interactive effects of time of day, pre and post paradigm, and group on plasma corticosterone (CORT) within a multilevel (mixed effects) model (*Equation 1*), with subject ID included as a random effect variable:
   *Equation 1: CORT ~* $\beta_0 + \beta_1(Stress) + \beta_2(TimeOfDay) + \beta_3(PreVsPost) + \beta_4(SIScore) + \beta_5(Stress*TimeOfDay)+$, *random=~1|ID*
2) Linear mixed-effects model fit by REML: CORT~ PreVsPostParadigm* TimeOfDay * Group
3) PreVsPostParadigm and TimeOfDay were set as dummy variables
4) Data distributions were examined for normality. SI Ratio was centered. Only CORT data required transformation due to non-normal skewed distribution—the data was log2 transformed and centered.
5) Time of day AM, Pre-Paradigm, and Control mice were set as the reference (intercept)

| Predictor | Estimate | Std.Error | DF | T.Value | P.Value |
|---|---|---|---|---|---|
| PreVsPostParadigm | 0.2912 | 0.5028 | 70 | 0.5792 | 0.5643 |
| **TimeOfDay** | **1.5521** | **0.5028** | **70** | **3.0868** | **0.0029** |
| GroupResilient | -0.4533 | 0.7111 | 33 | -0.6376 | 0.5482 |
| GroupSusceptible | -0.7169 | 0.6806 | 33 | -0.6126 | 0.0532 |
| PreVsPostParadigm:TimeOfDay | -0.4102 | 0.7055 | 70 | -0.5815 | 0.5628 |
| PreVsPostParadigm:GroupResilient | 0.1104 | 1.0056 | 70 | 0.1098 | 0.9128 |
| PreVsPostParadigm:GroupSusceptible | 0.8971 | 0.8369 | 70 | 1.0718 | 0.2875 |
| PreVsPostParadigm:TimeOfDay:GroupResilient | -0.3730 | 0.9737 | 70 | -0.3830 | 0.7028 |
| PreVsPostParadigm:TimeOfDay:GroupSusceptible | 1.1639 | 0.8210 | 70 | 1.41760 | 0.1607 |

### FIGURE 3 STATISTICS:

**Table S5: Coping Behaviors 2x2 Mixed ANOVA Results Table**
Behavior <- aov_ez("AnimalID", "SIRatio", Dataset, between = "Group", within = "Day", covariate=”Cohort”, type = "III")

| Behavior | Effect | F(df) | P.Value | Partial Eta Squared |
|---|---|---|---|---|
| ***Fight Duration*** | | | | |
| | Group | 0.03(1,44) | 0.860 | <0.001 |
| | Cohort | 1.34(2,44) | 0.272 | 0.027 |
| | **Day** | 16.20(1,44) | **<0.001** | 0.166 |
| | Group:Day | 0.19(1,44) | 0.665 | 0.002 |
| | Cohort:Day | 0(2,44) | >0.999 | <0.001 |
| ***Fight*** | | | | |
| | Group | 3.45(1,44) | 0.070 | 0.054 |
| | Cohort | 0.13(2,44) | 0.876 | 0.004 |
| | **Day** | 39.07(1,44) | **<0.001** | 0.266 |
| | Group:Day | 0(1,44) | 0.956 | <0.001 |
| | Cohort:Day | 3.05(2,44) | 0.057 | 0.054 |
| ***Escape*** | | | | |
| | **Group** | 7.07(1,44) | **0.011** | 0.092 |
| | Cohort | 1.34(1,44) | 0.193 | 0.047 |
| | **Day** | 29.92(2,44) | **<0.001** | 0.199 |
| | Group:Day | 1.09(1,44) | 0.303 | 0.009 |
| | Cohort:Day | 6.07(2,44) | 0.057 | 0.072 |
| ***Upright*** | | | | |
| | Group | 1.78(1,44) | 0.189 | 0.020 |
| | Cohort | 1.79(2,44) | 0.178 | 0.039 |
| | Day | 1.94(1,44) | 0.170 | 0.022 |
| | Group:Day | 0.95(1,44) | 0.335 | 0.011 |
| | Cohort:Day | 1.49(2,44) | 0.237 | 0.033 |
| ***Forward**** | | | | |
| | Group | 0.34(1,44) | 0.561 | 0.004 |
| | **Cohort** | 4.32(2,44) | **0.019** | 0.081 |
| | Day | 0.90(1,44) | 0.349 | 0.001 |
| | Group:Day | 0.01(1,44) | 0.917 | <0.001 |
| | **Cohort:Day** | 3.49(2,44) | **0.039** | 0.080 |
| ***Crouch**** | | | | |
| | Group | 0.46(1,44) | 0.502 | 0.003 |
| | **Cohort** | 4.44(2,44) | **0.018** | 0.064 |
| | Day | 2.44(1,44) | 0.125 | 0.035 |
| | Group:Day | 0.33(1,44) | 0.571 | 0.005 |
| | Cohort:Day | 2.18(2,44) | 0.125 | 0.062 |

*Forward and Crouch behaviors on average were utilized <5% and <7% of the time overall, therefore the variability as shown by the cohort effect is most likely due to the extremely low occurrences
-No Interaction effects were found, thus post-hoc tests were not conducted
-Coping Behavior correlations in Tables S14 and S15

### STATISTICS FIGURES 4 & 5:

#### Table S6: One-Way ANOVA Table

AnovaModel<-aov(OutputValue ~ Group)

| Behavior | F(df) | P.Value | Effect Size |
|---|---|---|---|
| **Open Field** | | | |
| *% Time Spent in Center (Group Effect)* | 6.689(2,31) | **0.0039** | 0.51 |
| *Total Distance Traveled (Group Effect)* | 1.718(2,31) | 0.1965 | 0.23 |
| **Forced Swim Test** | | | |
| *Time Spent Immobile (sec) (Group Effect)* | 3.93(2,31) | **0.0298** | 0.38 |
| **Von Frey** | | | |
| *Force of filament used (g) (Group Effect)* | 5.163(2,33) | **0.0112** | 0.44 |
| **Forced Interaction** | | | |
| *CORT (ng/mL) with CD1 Only (Group Effect)* | 4.56(2,23) | **0.0214** | 0.47 |
| *CORT (ng/mL) All Conditions (Group Effect)* | 11.44(2,32) | **0.0002** | 0.62 |
| **Body Weight** | | | |
| *Change in Body Weight Day 10 - Day (Group Effect)* | 18.6(2,33) | **<0.0001** | 0.71 |

#### Table S7: Linear Model for FIT CORT differences in Stressed Mice

FITCORT_defeat<-lm(FITCORT~SIRatio+BodyWeightChange)

| Effect | Estimate | Std.Error | T.Value | P.Value |
|---|---|---|---|---|
| **SIRatio** | **125.36** | **45.33** | **2.765** | **0.01711** |
| BodyWeightChange | -31.62 | 24.05 | -1.319 | 0.21170 |

#### Table S8: Post-Hoc Tukey Test Table

PostHocResults<- emmeans(dataset$anovamodel, ~AnimalID|Group, adjust="Tukey")

| Behavior | Group Comparison | P.adj |
|---|---|---|
| ***Open Field % Time Spent in Center*** | | |
| | Control-Resilient | **0.0074** |
| | Control-Susceptible | **0.0451** |
| | Resilient-Susceptible | 0.8648 |
| ***Forced Swim Test- Time Spent Immobile*** | | |
| | Control-Resilient | 0.1847 |
| | Control-Susceptible | 0.2712 |
| | Resilient-Susceptible | **0.0231** |
| ***Von Frey Test*** | | |
| | Control-Resilient | 0.8496 |
| | Control-Susceptible | **0.0087** |
| | Susceptible-Resilient | 0.0691 |
| **Forced Interaction Test (CD1 CORT)** | | |
| | Control - Resilient | **0.0162** |
| | Control - Susceptible | 0.4616 |
| | Susceptible - Resilient | 0.1413 |
| **Forced Interaction Test (overall CORT)** | | |
| | Control C57BL6J - Control CD1 | 0.2301 |
| | Control C57BL6J - Defeat CD1 | **0.0211** |
| | Control CD1 - Defeat CD1 | **0.0002** |
| ***Body Weight Change Day 10-Day 1*** | | |
| | Control-Resilient | 0.4574 |
| | Control-Susceptible | **<0.0001** |
| | Resilient-Susceptible | **0.0003** |

#### Table S9: 2x2 ANOVA Table Fig S3

MixedModel<-aov_ez("AnimalID", "Value", dataset, between="Group", within="Day", type= "III")

| Behavior | Effect | F(df) | P.Value | Partial eta squared |
|---|---|---|---|---|
| **Raw Body Weight Data** | | | | |
| | Group | 0.6(2,32) | 0.553 | 0.033 |
| | **Day** | 14.52(1,32) | **<0.001** | 0.037 |
| | **Group:Day** | 12.31(2,32) | **<0.001** | 0.062 |

### STATISTICS FIGURES 4 & 5 CONTINUED:

**Table S10: Pearson Correlations Table**
cor.test(dataset$Variable1, dataset$Variable2, method=Pearson)

| Behavior | Comparison | Pearson r | P.Value |
|---|---|---|---|
| **Open Field** | | | |
| | Control SI Ratio vs. % Time Spent in Center | 0.381 | 0.1701 |
| | Defeat SI Ratio vs. % Time Spent in Center | -0.172 | 0.5393 |
| **Forced Swim Test** | | | |
| | Control SI Ratio vs. Time Spent Immobile (sec) | 0.417 | 0.0757 |
| | Defeat SI Ratio vs. Time Spent Immobile (sec) | -0.264 | 0.3413 |
| **Von Frey Test** | | | |
| | Control Force (g) vs. SI Ratio | -0.132 | 0.6139 |
| | **Defeat Force (g) vs. SI Ratio** | **0.581** | **0.0144** |
| **Forced Interaction Test** | | | |
| | Control CORT ng/ml vs. SI Ratio (CD1) | 0.191 | 0.5963 |
| | Control CORT ng/ml vs. SI Ratio (C57BL6J) | -0.036 | 0.9195 |
| | **Defeat CORT ng/ml vs. SI Ratio** | 0.535 | **0.0260** |
| **Body Weight Changes** | | | |
| | Control SI Ratio vs. Body Weight (g) | 0.693 | 0.0943 |
| | **Defeat SI Ratio vs. Body Weight (g)** | 0.702 | **0.0035** |

### STATISTICS FIGURE 6:

**Table S11: 2x2 Mixed ANOVA Table**
MixedModel<-aov_ez(“AnimalID”, “Value”, dataset, between=”Group”, within=”Strain”, type= “III”)

| Behavior | Effect | F(df) | P.Value | Partial eta squared |
|---|---|---|---|---|
| **Novel C57BL6J Social Interaction** | | | | |
| | **Group** | 13.70(2,23) | **<0.001** | 0.443 |
| | Strain | 2.360(1,23) | 0.1380 | 0.033 |
| | **Group:Strain** | 18.11(2,23) | **<0.001** | 0.344 |
| **Black Swiss Social Interaction** | | | | |
| | **Group** | 13.270(2,20) | **<0.001** | .350 |
| | Strain | 0.790(1,20) | 0.3860 | 0.023 |
| | **Group:Strain** | 11.240(2,20) | **<0.001** | 0.400 |

**Table S12: One-Way ANOVA Table**
AnovaModel<-aov(SocialPartnerStrain ~ Group)

| Behavior | F(df) | P.Value | Effect Size |
|---|---|---|---|
| **Strain Specificity: C57BL6/J** | | | |
| *Social Group (Group Effect)* | 0.373(2,23) | 0.6930 | 0.17 |
| **Strain Specificity: CD1 Condition of C57BL6/J** | | | |
| *Social Group (Group Effect)* | 34.080(2,23) | **<0.0001** | 0.85 |
| **Strain Specificity: Black Swiss** | | | |
| *Social Group (Group Effect)* | 0.120(2,20) | 0.8880 | 0.11 |
| **Strain Specificity: CD1 Condition of Black Swiss** | | | |
| *Social Group (Group Effect)* | 19.390(2,20) | **<0.0001** | 0.79 |

**Table S13: Post-Hoc Tukey Tests**
PostHocResults<- emmeans(dataset$anovamodel, ~AnimalID|Group, adjust=”Tukey”)

| Behavior | Group Comparison | P.adj |
|---|---|---|
| ***Strain Specificity: C56BL6/J Interaction*** | | |
| | Control - Resilient | 0.8666 |
| | Control - Susceptible | 0.9294 |
| | Resilient - Susceptible | 0.6764 |
| ***Strain Specificity: CD1 Condition of C56BL6/J Interaction*** | | |
| | Control - Resilient | 0.4912 |
| | Control - Susceptible | **<0.0001** |
| | Resilient - Susceptible | **<0.0001** |
| ***Strain Specificity: Black Swiss*** | | |
| | Control - Resilient | 0.9153 |
| | Control - Susceptible | 0.8998 |
| | Resilient - Susceptible | 0.9956 |
| ***Strain Specificity: CD1 Condition of Black Swiss*** | | |
| | Control - Resilient | **0.0346** |
| | Control - Susceptible | **0.0000** |
| | Resilient - Susceptible | **0.0049** |

### Table S14: Pearson r Values for Coping Mechanism Correlations:

| | FightDay1 | EscapeDay1 | CrouchDay1 | ForwardDay1 | UprightDay1 | FightDurationDay1 | FightDay10 | EscapeDay10 | CrouchDay10 | ForwardDay10 | UprightDay10 | FightDurationDay10 | SIScore |
|---|---|---|---|---|---|---|---|---|---|---|---|---|---|
| **FightDay1** | 1 | **-0.511** | -0.159 | -0.278 | **-0.346** | **0.335** | 0.153 | -0.267 | 0.032 | 0.137 | 0.109 | 0.04 | **0.312** |
| **EscapeDay1** | | 1 | -0.187 | -0.122 | **-0.501** | -0.276 | 0.008 | 0.281 | -0.054 | **-0.372** | -0.196 | 0.084 | **-0.297** |
| **CrouchDay1** | | | 1 | **0.339** | -0.096 | 0.096 | 0.087 | -0.125 | -0.287 | 0.098 | 0.164 | 0.143 | -0.104 |
| **ForwardDay1** | | | | 1 | -0.001 | -0.125 | -0.166 | 0.182 | 0.01 | -0.096 | 0.007 | -0.079 | -0.107 |
| **UprightDay1** | | | | | 1 | -0.04 | -0.157 | -0.047 | 0.134 | 0.278 | 0.047 | -0.171 | 0.08 |
| **FightDurationDay1** | | | | | | 1 | -0.006 | -0.076 | 0.01 | 0.204 | -0.015 | -0.059 | 0.183 |
| **FightDay10** | | | | | | | 1 | -0.736 | -0.016 | -0.143 | -0.149 | **0.612** | -0.074 |
| **EscapeDay10** | | | | | | | | 1 | -0.208 | -0.157 | **-0.377** | **-0.513** | -0.088 |
| **CrouchDay10** | | | | | | | | | 1 | -0.186 | -0.008 | 0.088 | 0.059 |
| **ForwardDay10** | | | | | | | | | | 1 | -0.148 | -0.164 | -0.054 |
| **UprightDay10** | | | | | | | | | | | 1 | 0.023 | 0.277 |
| **FightDurationDay10** | | | | | | | | | | | | 1 | -0.126 |
| **SIScore** | | | | | | | | | | | | | 1 |

### Table S15 : P.Values for Coping Mechanism Correlations:

| | FightDay1 | EscapeDay1 | CrouchDay1 | ForwardDay1 | UprightDay1 | FightDurationDay1 | FightDay10 | EscapeDay10 | CrouchDay10 | ForwardDay10 | UprightDay10 | FightDurationDay10 | SIScore |
|---|---|---|---|---|---|---|---|---|---|---|---|---|---|
| **FightDay1** | NA | **0** | 0.281 | 0.056 | **0.016** | **0.020** | 0.299 | 0.067 | 0.83 | 0.352 | 0.462 | 0.787 | **0.031** |
| **EscapeDay1** | | NA | 0.203 | 0.407 | **0** | 0.058 | 0.957 | 0.053 | 0.718 | **0.009** | 0.181 | 0.572 | **0.04** |
| **CrouchDay1** | | | NA | **0.018** | 0.515 | 0.518 | 0.559 | 0.398 | 0.048 | 0.508 | 0.266 | 0.334 | 0.484 |
| **ForwardDay1** | | | | NA | 0.997 | 0.396 | 0.258 | 0.216 | 0.947 | 0.515 | 0.963 | 0.595 | 0.468 |
| **UprightDay1** | | | | | NA | 0.786 | 0.288 | 0.750 | 0.365 | 0.056 | 0.753 | 0.246 | 0.59 |
| **FightDurationDay1** | | | | | | NA | 0.965 | 0.610 | 0.948 | 0.163 | 0.918 | 0.690 | 0.214 |
| **FightDay10** | | | | | | | NA | 0 | 0.916 | 0.331 | 0.311 | **0** | 0.617 |
| **EscapeDay10** | | | | | | | | NA | 0.155 | 0.286 | **0.008** | **0** | 0.55 |
| **CrouchDay10** | | | | | | | | | NA | 0.206 | 0.956 | 0.553 | 0.691 |
| **ForwardDay10** | | | | | | | | | | NA | 0.316 | 0.265 | 0.715 |
| **UprightDay10** | | | | | | | | | | | NA | 0.875 | 0.057 |
| **FightDurationDay10** | | | | | | | | | | | | NA | 0.392 |
| **SIScore** | | | | | | | | | | | | | NA |
